# Historical contingency is revealed by trade-offs upon sequential adaptation to different infection routes

**DOI:** 10.64898/2026.11.05.764508

**Authors:** Priscilla A. Akyaw, Élio Sucena, Elvira Lafuente

**Author notes:** Equal contributions (Élio Sucena, Elvira Lafuente). Current address: Institute of Ecology and Evolution, School of Biological Sciences, University of Edinburgh, Edinburgh, United Kingdom (Priscilla A. Akyaw).

## Abstract

Past adaptation can shape how populations respond to subsequent selection. Because host-pathogen interactions vary widely in their ecological and physiological context, they provide a powerful system for testing the role of historical contingency in adaptation. Here, we evolved replicate populations of *Drosophila melanogaster*, previously adapted to systemic infection with *Pseudomonas entomophila*, under oral exposure to the same pathogen. Both evolutionary histories evolved increased survival under oral infection, but previous systemic adaptation altered the trajectory of this response in a sex-dependent manner. Females with a history of systemic selection began with lower oral-infection survival, yet adapted more rapidly than controls, whereas males showed weaker history-dependent differences. These contrasting trajectories could not be attributed simply to an overall loss of standing genetic variation in the founder populations. Adaptation to oral infection also incurred a trade-off, reducing survival under systemic infection, whereas the fecundity cost of infection did not differ between evolutionary histories. Finally, sex- and generation-specific changes in bacterial clearance revealed that adaptation to oral infection involved distinct mechanistic responses alongside shared genetic components. Together, our results show that past selection shapes both the tempo and mechanism of adaptation, revealing that responses to different infection ecologies are built on partially shared, but also route-specific, genetic architectures.

## INTRODUCTION

Populations adapting to similar environments may evolve similar phenotypic outcomes while relying on markedly different genetic changes and physiological mechanisms. This variation reflects historical contingency: past selection, genetic drift, and evolutionary constraints have shaped present genetic architecture and, consequently, define the adaptive potential of the population (Bull & Charnov, 1985; Gould, 1989; Travisano *et al*., 1995). Classic experimental and comparative studies have shown that even small past differences can produce divergent trajectories under similar selective pressures (Losos, Warheitt & Schoener, 1997; Blount, Borland & Lenski, 2008; Meyer *et al*., 2012). Understanding how historical contingency affects adaptation is key to predicting both the repeatability of evolution and the diversification of lineages (Teotónio & Rose, 2001, 2004; Stern, 2013; Fragata *et al*., 2014; Matos *et al*., 2015; Lenski, 2017). For example, prior adaptation may enhance performance in related environments through pleiotropic effects or the reuse of shared physiological pathways (Futuyma & Moreno, 1988). Conversely, adaptation to one environment may favor specialization and constrain future evolution because directional selection erodes standing genetic variation (Levins, 1968; Fry, 1996; Hereford, 2009) or through trade-offs, whereby beneficial changes in one context are costly in another (Stearns, 1992; Roff, 2002).

In the context of immunity, such trade-offs arise from the energetic and physiological costs of mounting and maintaining immune defenses (Lochmiller & Deerenberg, 2000). Ecological immunology has provided abundant evidence for these costs, which can reduce fecundity, lifespan, or competitive ability across taxa (Sheldon & Verhulst, 1996; Lochmiller & Deerenberg, 2000; Schmid-Hempel & Ebert, 2003; McKean *et al*., 2008). In *Drosophila melanogaster*, immune activation has repeatedly been shown to impose reproductive and metabolic costs (Flatt, 2011; Faria *et al*., 2015; Fabian *et al*., 2018; Singh *et al*., 2025). Such physiological constraints create the potential for evolutionary trade-offs between infection resistance and other life-history traits, linking proximate mechanisms to long-term evolutionary trajectories.

*Drosophila* are exposed to pathogens through several routes. Although systemic infection can occur through wounding, oral ingestion is considered one of the most common infection routes in nature (Vodovar *et al*., 2005; Siva-Jothy *et al*., 2018; Subasi *et al*., 2024). Importantly, systemic and oral infections engage overlapping but distinct defense mechanisms (Martins *et al*., 2013; Mondotte & Saleh, 2018). Systemic infection bypasses epithelial barriers and activates humoral and cellular immune responses in the hemocoel, whereas oral infection relies strongly on gut epithelial defenses, local immune signaling, digestive physiology, and interactions with the microbiota (Buchon *et al*., 2009; Kounatidis & Ligoxygakis, 2012; Broderick & Lemaitre, 2012; Akyaw *et al*., 2025). Consistent with this mechanistic distinction, experimental evolution studies have found limited evidence for correlated responses or trade-offs between adaptive outcomes to systemic and oral infections (Faria *et al*., 2015; Gupta *et al*., 2016, 2017; Ahlawat *et al*., 2022; Paulo *et al*., 2025). These findings suggest that the physiological systems underlying defense against each route are at least partly independent. However, whether this mechanistic distinction makes adaptation to different infection routes evolutionarily independent remains unresolved.

In an earlier work, *D. melanogaster* populations selected under systemic infection with *Pseudomonas entomophila* evolved increased survival relative to control populations exposed only to injury (Martins *et al*., 2013). These populations provided an opportunity to test whether prior adaptation to one infection route shapes subsequent adaptation to another route with the same pathogen. Here, we subjected populations with a history of systemic infection selection (hereafter BactSys) and matched control histories (hereafter ContSys) to experimental evolution under oral *P. entomophila* infection, generating BactSysOral and ContSysOral populations. Across the experiment, we tracked survival, bacterial clearance, systemic-infection performance, and fecundity to ask whether historical background influences the tempo and direction of adaptation to oral-infection.

## MATERIALS AND METHODS

### Maintenance of *Drosophila* populations and experimental evolution under oral selection

The experimental histories used here originated from a previous selection experiment in which BactSys populations evolved under systemic infection by pricking flies with pathogen bacteria *Pseudomonas entomophila*, whereas ContSys populations experienced the same handling regime without live bacterial systemic infection (i.e. pricking with sterile PBS) (Martins *et al*., 2013, 2014).

At the start of the present experiment, each history was split into four replicate populations, generating four BactSys-derived and four ContSys-derived replicate lineages. These eight replicate populations were then subjected to selection under oral infection with *P. entomophila* and are referred to as BactSysOral and ContSysOral after oral selection. For each generation of oral selection, 320 females and 320 males per replicate population were collected as 3–5-day-old adults and exposed to oral infection with *P. entomophila*. After infection, flies were returned to their respective cages with clean food and allowed to recover and mate for six days. Only individuals surviving infection after the six-day period contributed to egg laying and therefore founded the next generation. Selection was maintained until generation 26, after which populations were kept under relaxed selection and analyzed separately where indicated. Oral selection was conducted every generation and survival was assayed after infection in each generation (see ‘Survival and fecundity assays’ section) except G2, G14, and G21, when phenotyping could not be performed because of technical constraints.

All replicate populations were maintained at 25°C, 55–70% relative humidity, and a 12:12 h light:dark photoperiod on standard cornmeal-molasses-agar medium. Generations were discrete and lasted approximately three weeks. Controlled egg lays were used to standardize larval density across replicate populations and generations.

### Pathogen culture and infection protocols

Oral infections used a rifampicin-resistant *P. entomophila* strain, a generous contribution from the Lemaitre lab (Vodovar et al., 2005). Bacteria were streaked on LB agar, and a single colony was used to initiate an overnight culture in LB medium at 29°C with shaking. Cells were collected by centrifugation, washed, resuspended in sterile PBS, adjusted to OD600 = 100, and mixed 1:1 with 5% sucrose. For oral-infection assays, adult flies were briefly anesthetized with CO₂, sorted by sex into vials, allowed to recover for at least 24 h, and then transferred to vials containing filter paper saturated with the bacterial sucrose suspension. Control flies experienced the same handling and exposure conditions but received 5% sucrose without live bacteria. After exposure, flies were transferred to fresh food and monitored for survival or used for downstream phenotyping as described below.

Systemic infection was assayed in ancestral BactSys and ContSys populations and in BactSysOral and ContSysOral populations after oral selection. Flies were sorted by sex into vials using the sex-specific starting numbers recorded in the raw backbone file. Flies were pricked in the thorax with a fine needle dipped into *P. entomophila* adjusted to OD_600_ = 0.01. Sham-infected controls were pricked with PBS. Mortality was recorded over the following days.

### Survival and fecundity assays

In both oral- and systemic-infection survival assays, mortality was recorded separately for females and males for six days. Oral selection continued every generation through G26, but survival phenotyping could not be performed at G2, G14, and G21 because of technical constraints; these generations are therefore absent from the selected-trajectory analyses.

Fecundity was analyzed for selected generations G4, G9, and G23. Fecundity data from ancestral and relaxed-selection assays were analyzed separately. Vial-level mortality-corrected fecundity was calculated by first dividing the number of pupae counted each day by the number of females alive on the preceding day, then summing these daily pupae-per-alive-female values across the assay. To focus on the infection-specific reproductive response and account for baseline differences among generations, blocks, and populations, fecundity under PeWT was normalized to fecundity under sucrose controls within each population and replicate. Values below 1 therefore indicate reduced fecundity under infection relative to the matched sucrose control.

### Bacterial load assays

Bacterial clearance after live oral infection was assayed at G4, G8, and G16 using previously established protocols with modifications (Nehme & Poole, 2007; Martins *et al*., 2013; Neyen *et al*., 2014). Briefly, flies collected at the designated sampling time points after specific time points after infection were surface-sterilized with 70% ethanol, followed by 60% bleach and two washes with Milli-Q water. Sterilized flies were subsequently placed in 96-deep-well plates, where they were homogenized with glass beads in 50 µl of sterile LB using a TissueLyser (Qiagen). Homogenates were serially diluted through five successive 1:10 dilution steps, after which 4 µl of each dilution was plated on LB agar supplemented with rifampicin. Colonies were allowed to grow overnight at 29°C. The following day, all plates were scanned, and colonies were counted from digital images.

### DNA extraction and pool-sequencing

Genomic DNA was extracted using a salt–chloroform protocol adapted from Miller, Dykes & Polesky (1988). For each pool, approximately 120–130 adult flies were placed in a 50-ml tube with six to eight beads, homogenized by vortexing in 5 ml of homogenization buffer (160 mM sucrose, 80 mM EDTA, pH 8.0, 100 mM Tris, pH 8.0, and 0.5% SDS) supplemented immediately before use with proteinase K to a final concentration of 0.10 mg/ml, and incubated overnight at 58 °C (minimum 4–6 h). A 200-µl aliquot of each lysate was then treated with 2 µl RNase A (10 mg/ml) for 30–60 min at room temperature. Proteins and cellular debris were removed by adding 100 µl of 4.5 M NaCl and extracting with 225 µl chloroform for 10 min at room temperature, followed by centrifugation for 10 min at maximum speed at 4 °C. The upper aqueous phase (approximately 200 µl) was transferred to a new tube, avoiding the interphase, and DNA was precipitated with an equal volume of 100% isopropanol. After centrifugation for 10 min at maximum speed, the pellet was washed with 0.5 ml of 70% ethanol, incubated for 15 min at room temperature, centrifuged again for 10 min, air-dried, and resuspended in 50 µl of double-distilled water (ddH₂O).

At G0, four replicate DNA pools were prepared from each selection history (BactSys and ContSys). Because the sex composition of the pools was not recorded, analyses assumed an equal sex ratio. Sequencing reads were aligned to the *Drosophila melanogaster* reference genome (BDGP6.32) prior to downstream analyses. Analyses were restricted to chromosomes 2L, 2R, 3L, 3R, and X. Allele counts were obtained using samtools mpileup, thereby avoiding genotype calls based on a diploid model. Reads and bases with mapping or base quality below 20 were excluded, overlapping mates were counted once, BAQ was disabled, and indel strings were omitted. Candidate sites required a total non-reference count of at least eight across the eight G0 pools, including at least two forward- and two reverse-strand observations. Analyses retained biallelic sites with a read depth of 10–150 in every pool, a global minor-allele count of at least eight, and a third-allele fraction no greater than 2%.

### Population-genetic analyses

Pairwise differentiation was estimated from synchronized pooled read counts using the Karlsson estimator implemented in PoPoolation2 (Kofler, Pandey & Schlötterer, 2011). Sites were retained when coverage was 10–150 reads in every pool and the pooled minor-allele count was at least eight. FST was calculated in non-overlapping 50-kb windows, with windows lacking informative SNPs suppressed, and each contrast was summarized as the mean window estimate weighted by the number of SNPs in each window. As an independent robustness check, we also calculated a Hudson-type ratio-of-sums estimator from the pooled read counts. For this check, estimates were corrected for finite biological-pool sampling and repeated assuming 120, 125, or 130 flies per pool; autosomes were assigned 2N sampled chromosomes and the X chromosome 1.5N under the assumed equal sex ratio.

Expected heterozygosity was calculated for each replicate pool across the common filtered polymorphic-site set using an unbiased read-count estimator with the same finite-pool correction. The midpoint assumption of 125 flies was used for heterozygosity reporting, and the BactSys–ContSys difference was evaluated using a chromosome-stratified bootstrap of paired non-overlapping 50-kb windows (5,000 replicates). Because invariant callable sites were not included, these heterozygosity values describe diversity within the shared ascertained set of polymorphic sites.

### Statistical analyses

All analyses were performed in R. Data processing and visualization used tidyverse and ggplot2. Oral- and systemic-infection Cox mixed models used coxme, estimated marginal means and trends were obtained using emmeans, and count or binomial mixed models used glmmTMB. Model diagnostics were assessed using simulation-based residual checks where applicable.

Oral-infection survival across the selected trajectory was analyzed using Cox mixed-effects models. Models were fitted separately by sex and treatment. The main continuous-generation model included Population, Generation, and their interaction as fixed effects, with replicate population included as a random intercept to account for baseline differences among replicate evolutionary lineages. To test whether the pace of adaptation changed over time, we fitted segmented Cox mixed-effects models with separate early and late generation slopes and compared one-slope and two-phase models across candidate breakpoints; the best-supported breakpoint occurred at G5.

Generation-specific BactSysOral *versus* ContSysOral contrasts were calculated from the continuous-generation Cox mixed model evaluated at each observed generation. For Cox model fitting only, all-censored vials were handled by coding one individual at the final observation time as an event. The observed event column was retained unchanged for raw survival summaries and plots.

Fecundity analyses used the normalized PeWT/sucrose fecundity ratio as the response. Ratios were log-transformed and analyzed with linear models including population or selection history, stage/generation, and their interaction. Separate models were fitted for the selected trajectory and for the ancestral/relaxed-selection assays. Estimated marginal means and population contrasts were back-transformed to ratios for reporting.

Bacterial-clearance analyses first modeled raw colony counts across serial dilutions using negative-binomial generalized linear mixed models. For each generation, counts were modeled as a function of log10 dilution, continuous time after infection, population, sex, and their interactions, with fly identity included as a random effect allowing dilution-specific variation and replicate population included as an additional random effect. High-order dilution interactions were simplified before estimating clearance rates. Clearance slopes were estimated with emtrends for each population and sex at a reference dilution, and pairwise contrasts tested whether BactSysOral and ContSysOral differed in clearance rate. Bacterial half-life was calculated as log(2) divided by the estimated clearance rate.

As a follow-up analysis, we fitted a two-part model to separate zero-CFU observations from bacterial-load dynamics among flies with detectable bacteria. First, fly-level detection was modeled as a binomial response indicating whether a fly had zero detectable CFU, with continuous time, population, sex, and their interactions as predictors and replicate population as a random effect. Second, among flies with positive CFU values, log10-transformed bacterial load was modeled with continuous time, population, sex, and their interactions, again including replicate population as a random effect. Population differences were summarized as differences in the time slope for the zero-CFU component and as differences in clearance rate for the positive-load component.

Systemic-infection survival was analyzed using Cox mixed-effects models. Models were fitted separately by sex and treatment to compare the four populations directly and to test Stage, Selection history, and their interaction, where Stage distinguished ancestral from after-selection populations and Selection history distinguished BactSys-derived from ContSys-derived histories. Vial identity was included as a random intercept to account for shared vial-level mortality risk. To test whether oral selection carried a cost for systemic-infection performance, an additional PeWT Cox mixed model tested after-selection *versus* ancestral populations within each selection history while accounting for sex. For visualization of endpoint survival, the number of flies surviving the final mortality check was analyzed with quasibinomial models using sex-specific vial starting counts as denominators, followed by estimated marginal means and pairwise contrasts.

## RESULTS

### Survival dynamics under oral infection

Survival after oral infection increased during experimental evolution in both populations and sexes, indicating rapid adaptation to the oral-infection regime (Fig. 1A; Table S4). This response was infection-specific: survival under the sucrose control remained high throughout the experiment and showed no comparable directional increase (Fig. S1).

**Figure 1.**
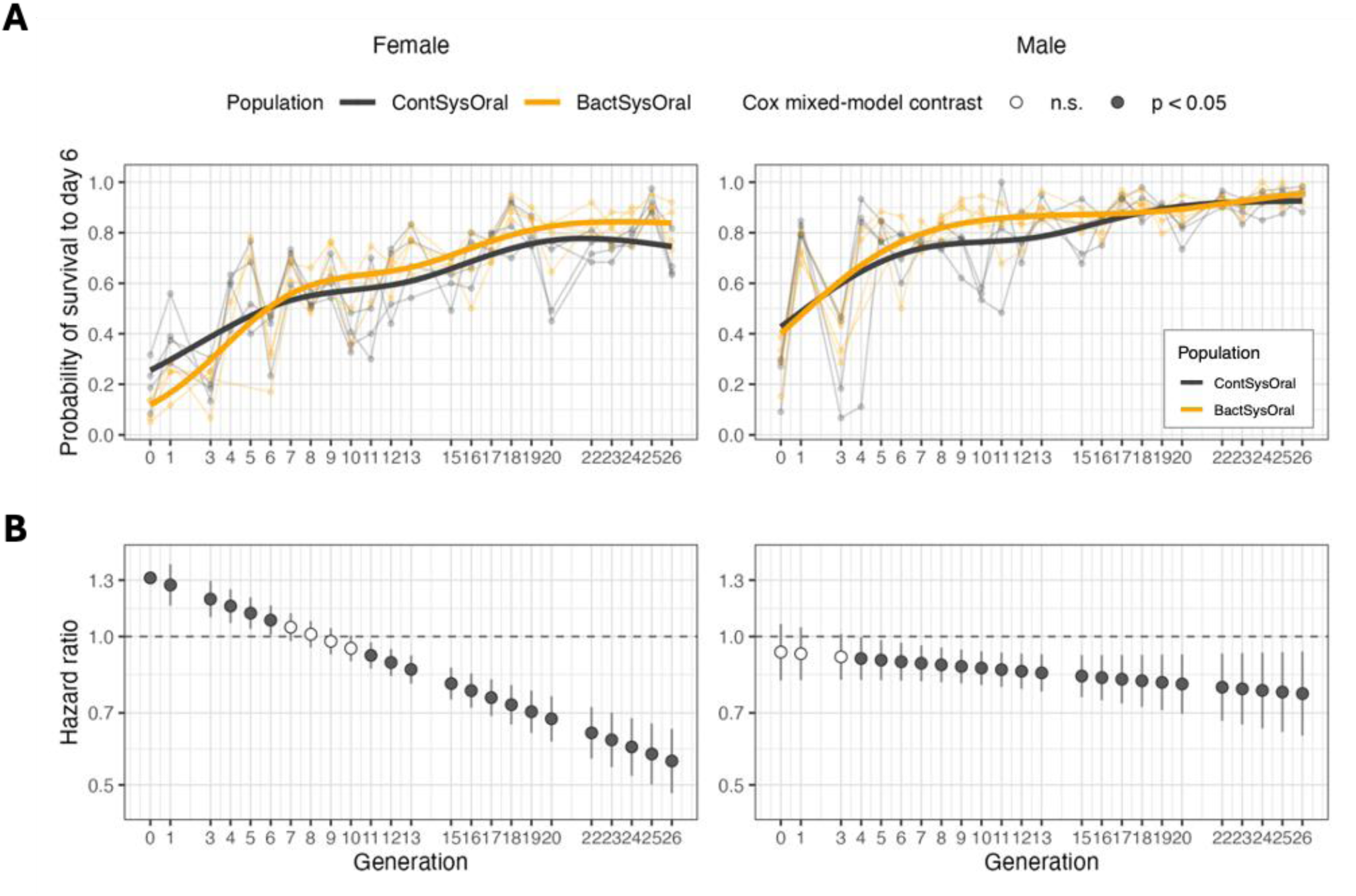
Survival adaptation during experimental evolution under oral infection. Survival of BactSysOral and ContSysOral populations across the selected trajectory. **A**. Probability of survival to six days after oral *Pseudomonas entomophila* infection across generations; thin lines show replicate populations, and thick lines show model-predicted population means. **B**. Generation-specific BactSysOral/ContSysOral hazard ratios from Cox mixed models; values below 1 indicate lower mortality in BactSysOral relative to ContSysOral, and filled points indicate p < 0.05. Female and male results are shown in separate columns. Gaps at G2, G14, and G21 indicate generations in which survival was not assayed because of technical constraints.

Survival increased with generation in ContSysOral females (Hazard ratio (HR) per generation = 0.93, 95% CI: 0.93–0.94, p < 0.001). This increase was stronger in BactSysOral females (Population x Generation HR ratio = 0.97, 95% CI: 0.96–0.98, p < 0.001), indicating faster adaptation in populations with a previous history of systemic infection. Males also evolved increased survival (ContSysOral HR per generation = 0.92, 95% CI: 0.91–0.92, p < 0.001). However, the BactSysOral *versus* ContSysOral difference in the overall linear slope was small and not significant in males (Population x Generation HR ratio = 0.99, 95% CI: 0.98–1.00, p = 0.18), suggesting that the male response was not well summarized by a single population-specific linear slope.

We therefore fitted segmented Cox mixed models to test whether adaptation changed pace over time. In both sexes, survival was better described by a two-phase model than by a single linear generation slope, with the best-supported breakpoint at G5 (females: delta AIC = 67.90, LRT chi-square = 71.85, p < 0.001; males: delta AIC = 85.30, LRT chi-square = 89.19, p < 0.001). In females, both populations improved rapidly before G5 (ContSysOral: HR per generation = 0.87, 95% CI: 0.84–0.89; BactSysOral: HR per generation = 0.82, 95% CI: 0.80–0.84; both p < 0.001). The early slope was steeper in BactSysOral than in ContSysOral females (slope HR ratio = 0.95, 95% CI: 0.91–0.99, p < 0.01), and BactSysOral females also improved faster after G5 (slope HR ratio = 0.98, 95% CI: 0.96–0.99, p < 0.001). In males, both populations again improved rapidly before G5 (ContSysOral: HR per generation = 0.83, 95% CI: 0.80– 0.86; BactSysOral: HR per generation = 0.79, 95% CI: 0.76–0.82; both p < 0.001). The BactSysOral *versus* ContSysOral early slope difference was marginal (slope HR ratio = 0.95, 95% CI: 0.91–1.00, p = 0.06), and late-phase slopes did not differ between populations (slope HR ratio = 1.00, 95% CI: 0.99–1.02, p = 0.64).

Generation-specific Cox mixed-model contrasts supported these sex-specific temporal patterns (Fig. 1B; Table S1). In females, BactSysOral initially had higher mortality than ContSysOral (G0 HR = 1.314, 95% CI: 1.185–1.457, p < 0.001), but this difference disappeared around intermediate generations and then reversed: BactSysOral females had lower mortality by G11 (HR = 0.915, 95% CI: 0.861–0.973, p < 0.01), with a stronger difference by G26 (HR = 0.559, 95% CI: 0.481–0.650, p < 0.001). In males, there was no detectable initial difference between populations (G0 HR = 0.929, 95% CI: 0.815–1.060, p = 0.28), but BactSysOral males had lower mortality by G4 (HR = 0.902, 95% CI: 0.816–0.997, p = 0.04) and remained lower at G26 (HR = 0.766, 95% CI: 0.629–0.932, p < 0.01). Complete generation-specific contrasts are reported in Table S1.

Thus, both histories evolved higher survival under oral infection, but prior systemic selection changed the path of this response. The effect depended on sex: BactSysOral females began with lower survival and then improved faster than ContSysOral females, whereas males showed a clear two-phase response with weaker population-specific slope differences.

We also compared ancestral BactSys and ContSys at G0 to evaluate their relative genetic composition. Using the Karlsson estimator, the SNP-weighted mean FST across informative non-overlapping 50-kb windows was 0.153 between histories, compared with 0.0046 among replicate pools within BactSys and 0.0048 within ContSys. An independently calculated Hudson-type ratio-of-sums estimate gave the same conclusion (between-history FST = 0.155; within-history FST = 0.0005 and 0.0007 for BactSys and ContSys, respectively). Mean expected heterozygosity across the common filtered polymorphic-site set was 0.246 in BactSys and 0.271 in ContSys, corresponding to approximately 9% lower heterozygosity in BactSys (BactSys − ContSys = −0.0256; chromosome-stratified 50-kb-window bootstrap 95% CI: −0.0279 to −0.0234). Thus, the populations entered oral selection with different allele-frequency compositions and with moderately lower diversity in BactSys.

### Bacterial Clearance Across Generations of Oral Infection

We then tested whether oral selection changed bacterial clearance after live *P. entomophila* infection (Table S5). Bacterial clearance was assayed at G4, G8, and G16 by quantifying bacterial loads over time after live oral infection. Clearance rates were compared between BactSysOral and ContSysOral populations within each sex; positive clearance-rate differences indicate faster bacterial clearance in BactSysOral relative to ContSysOral (Fig. 2).

**Figure 2.**
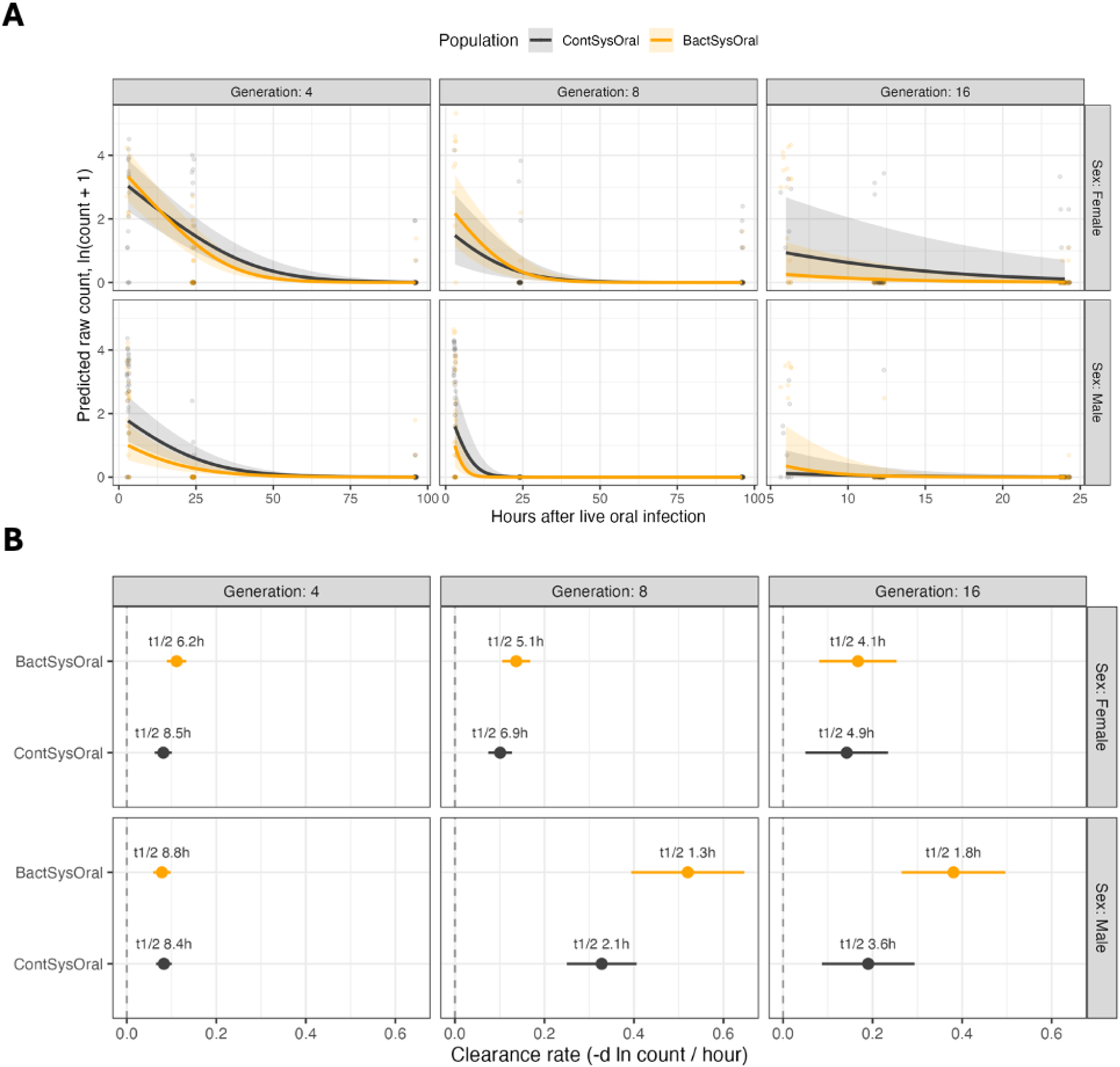
Bacterial clearance after live oral infection. Clearance of live *Pseudomonas entomophila* after oral infection at selected generations G4, G8, and G16. **A**. Model-based bacterial load trajectories over time, estimated from raw dilution-count models at the reference dilution. **B**. Estimated clearance rates by population, sex, and generation. Faster clearance corresponds to steeper declines in bacterial load and shorter estimated bacterial half-life. Points and intervals show model estimates with 95% confidence intervals.

At G4, BactSysOral females cleared bacteria faster than ContSysOral females in the raw dilution-count model (clearance-rate difference = 0.03, 95% CI: 0.00–0.06; p = 0.04). This corresponded to a shorter estimated bacterial half-life in BactSysOral females than in ContSysOral females (6.2 h *vs.* 8.5 h). In contrast, males showed no detectable population difference at G4 (clearance-rate difference near zero, 95% CI: −0.03 to 0.02; p = 0.73), with similar bacterial half-lives in BactSysOral and ContSysOral males (8.8 h vs. 8.4 h).

By G8, the female difference was weaker and no longer statistically significant, although the direction remained consistent with faster clearance in BactSysOral females (clearance-rate difference = 0.036, 95% CI: −0.004 to 0.075; p = 0.079; half-life: 5.1 h vs. 6.9 h). In males, however, BactSysOral flies cleared bacteria faster than ContSysOral flies (clearance-rate difference = 0.19, 95% CI: 0.05–0.34; p = 0.01), corresponding to bacterial half-lives of 1.3 h and 2.1 h, respectively.

At G16, BactSysOral females again showed only a weak and non-significant tendency toward faster clearance (clearance-rate difference = 0.03, 95% CI: −0.10 to 0.15; p = 0.68; half-life: 4.1 h *vs*. 4.9 h). In males, BactSysOral cleared bacteria faster than ContSysOral (clearance-rate difference = 0.19, 95% CI: 0.05–0.33; p < 0.01). The estimated bacterial half-life was approximately half as long in BactSysOral males as in ContSysOral males (1.8 h *vs.* 3.6 h).

Because zero-CFU observations could reflect complete clearance, low bacterial burden, or differences in ingestion, we followed the raw-count analysis with a two-part model. This model separated the probability of zero detectable CFU from bacterial-load decline among flies with detectable bacteria. The zero-CFU component showed no significant BactSysOral *versus* ContSysOral difference in the rate at which zero-CFU flies accumulated over time in any generation or sex (all 95% CIs overlapped zero; all p >= 0.12). In the positive-load component, only G8 males retained a significant BactSysOral advantage (clearance-rate difference = 0.11, 95% CI: 0.03–0.20; p < 0.01). Other positive-load contrasts were not significant (all p >= 0.06).

Together, these results indicate that oral-infection adaptation was accompanied by faster bacterial clearance in some contexts, but that clearance did not change uniformly across sexes or generations (Fig. 2).

### Systemic infection assays across generations

We also tested whether oral selection changed the response to systemic infection (Table S6). Survival upon systemic infection revealed a clear effect of selection history: BactSys-derived populations generally survived systemic *P. entomophila* wild type (PeWT) infection better than ContSys-derived populations, whereas PBS-pricked controls showed high survival and no detectable population differences (Fig. 3A; Fig. S2). In females, BactSys flies had lower mortality than ancestral ContSys flies following systemic PeWT infection in the Cox mixed model (HR = 0.39, 95% CI: 0.19–0.80, p < 0.01), and this difference was also evident after selection, with BactSysOral females surviving better than ContSysOral females (HR = 0.31, 95% CI: 0.17–0.58, p < 0.001). Endpoint survival estimates showed the same pattern: under PeWT, predicted survival at G46 was 0.71 in BactSys *versus* 0.39 in ContSys females, and 0.56 in BactSysOral *versus* 0.23 in ContSysOral females.

**Figure 3.**
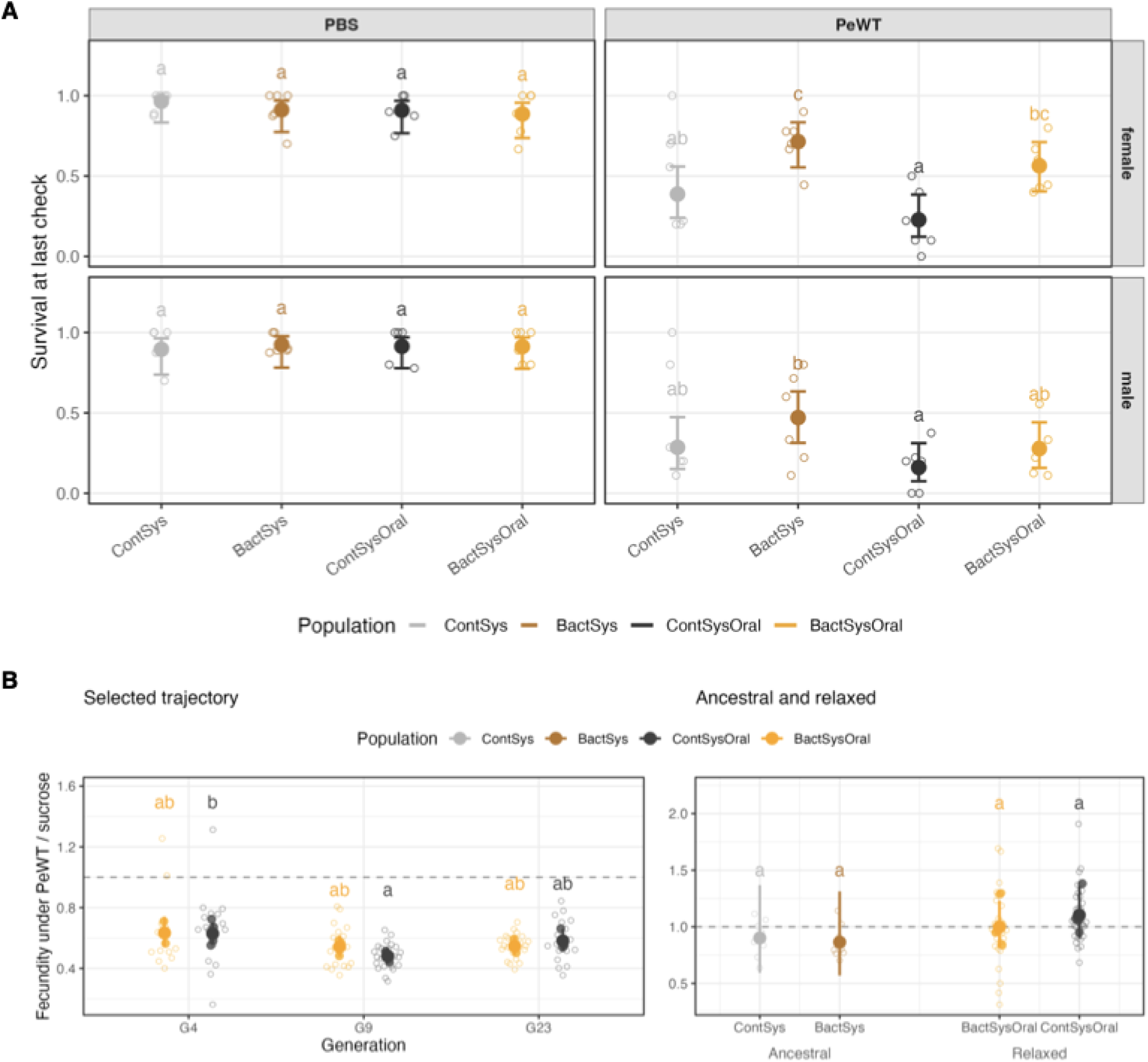
Systemic-infection survival and reproductive output after oral-infection adaptation. **A.** Survival at the last mortality check after systemic challenge with *Pseudomonas entomophila* or control PBS. Small open circles show vial-level survival, and large points with error bars show estimated marginal means with 95% confidence intervals. Letters indicate compact-letter groupings within each sex and treatment panel. Full survival curves are shown in Fig. S2. **B.** Fecundity under oral PeWT infection normalized to fecundity under sucrose controls. Values below 1 indicate reduced reproductive output under infection. Small open circles show replicate vials, filled smaller points show replicate population means where available, and large points with error bars show model-estimated marginal means with 95% confidence intervals. The selected trajectory includes G4, G9, and G23; ancestral and relaxed-selection assays are shown separately. Compact-letter groupings indicate post hoc comparisons within each panel.

In males, the ancestral BactSys *versus* ContSys difference was weaker and not statistically significant (Cox HR = 0.65, 95% CI: 0.36–1.17, p = 0.15), but after selection BactSysOral males had lower mortality than ContSysOral males (Cox HR = 0.57, 95% CI: 0.34–0.95, p = 0.03). Endpoint survival estimates followed the same direction, with predicted PeWT survival of 0.28 in BactSysOral males and 0.16 in ContSysOral males. Oral-selected populations also showed reduced systemic-infection survival relative to their corresponding ancestral population. This reduction was clear in the ContSys history (ContSysOral *versus* ContSys: Cox HR = 1.76, 95% CI: 1.16–2.67, p < 0.01) and weaker but in the same direction in the BactSys history (BactSysOral *versus* BactSys: Cox HR = 1.54, 95% CI: 0.97–2.44, p = 0.07). The magnitude of this cost was not detectably different between histories (Assay stage × Selection history interaction: HR = 0.87, 95% CI: 0.47–1.62, p = 0.67). Thus, BactSys-derived populations retained their advantage under systemic PeWT challenge, but oral selection was accompanied by reduced systemic-infection performance relative to the corresponding ancestral histories.

### Fecundity during adaptation to oral infection

Finally, we tested whether oral-infection adaptation was accompanied by changes in reproductive output (Table S7). Across the selected trajectory, infection reduced fecundity relative to sucrose controls at G4, G9 and G23, indicating a reproductive cost of oral infection (Fig. 3B). Across populations and generations, estimated PeWT/sucrose fecundity ratios ranged from 0.48 to 0.63 (all 95% confidence intervals excluded 1 and all p < 0.001). However, the magnitude of this infection-specific fecundity reduction did not differ significantly between BactSysOral and ContSysOral populations. The normalized PeWT/sucrose fecundity ratio was similar between populations at G4 (BactSysOral/ContSysOral ratio = 1.01, 95% CI: 0.83–1.22, p = 0.94), G9 (ratio = 1.13, 95% CI: 0.95–1.34, p = 0.15), and G23 (ratio = 0.94, 95% CI: 0.79–1.11, p = 0.43).

The ancestral and relaxed-selection assays allowed us to compare reproductive responses before oral selection and after subsequent relaxation. After normalization to sucrose controls, infection-specific fecundity did not differ significantly between ancestral BactSys and ContSys histories (BactSys/ContSys ratio = 0.96, 95% CI: 0.53– 1.73, p = 0.87). Similarly, after relaxed selection, BactSysOral and ContSysOral did not differ significantly in normalized infection fecundity (BactSysOral/ContSysOral ratio = 0.91, 95% CI: 0.67–1.22, p = 0.44).

Oral infection therefore reduced fecundity, but this reduction did not differ consistently between histories or after oral selection. Thus, the survival gains observed during adaptation were not mirrored by infection-specific reproductive output.

## DISCUSSION

### Past selection shaped adaptation to a different infection route

Historical contingency is usually discussed as a property of evolutionary trajectories: populations may face similar selection but respond differently because earlier selection has shaped the genetic and physiological contexts on which later selection acts (Bull & Charnov, 1985; Gould, 1989; Travisano *et al*., 1995). Survival after infection can reflect resistance, tolerance, tissue repair, behavior, metabolism, and life-history allocation, and these components need not evolve together (Schmid-Hempel, 2005; Lazzaro & Little, 2009). This mechanistic complexity which characterizes host-pathogen interactions sets a fruitful stage for testing historical contingency and its underlying genetic correlations. In our experiment, both BactSysOral and ContSysOral populations adapted to oral *Pseudomonas entomophila* infection, with the two histories differing both in their initial performance and in their subsequent trajectories under oral selection. This shows that adaptation to infection via different routes, an ecologically relevant circumstance, is genetically correlated and imposes relevant (historical) contingencies upon adaptation.

In females, the history of systemic selection initially carried a cost under oral infection: BactSysOral females survived poorly relative to ContSysOral females at the start of oral selection. However, this disadvantage was not a fixed constraint, as observed over the course of the oral selection. BactSysOral females improved faster, and later survived better than ContSysOral females. This pattern suggests that past systemic selection altered both the starting point and the tempo of adaptation to the new infection route. In males, the initial survival difference between histories was not evident, but survival was still better described by two phases of adaptation, and BactSysOral males later showed lower mortality than ContSysOral males. Thus, the legacy of past selection was sex-dependent: in females it affected both initial performance and subsequent adaptation rate, whereas in males it was expressed more clearly in the later outcome of oral selection.

Similarly to a previous study where an outbred *D. melanogaster* population (referred to as BactOral) adapted to oral infection with *P. entomophila* (Martins et al., 2013; Paulo et al., 2025), we observe here that improved survival after oral infection was rapid, with almost double the survival rate of both BactSysOral and ContSysOral after just five generations of selection. These results therefore conform with our previous work and that of others showing that *Drosophila* does evolve rapidly in response to different pathogenic bacteria (Ye, Chenoweth & McGraw, 2009; Gupta *et al*., 2016; Geeta Arun *et al*., 2021; Ahlawat *et al*., 2022).

At G0, genetic differentiation between BactSys and ContSys was substantial (SNP-weighted mean window FST = 0.153), whereas differentiation among replicates within each history was much lower (0.0046 in BactSys and 0.0048 in ContSys). These genomic results indicate that previous selection had driven genetic divergence between the two populations, consistent with previously reported phenotypic differences (Martins et al. 2013). In addition, heterozygosity was 9% lower in BactSys than in ContSys across the common set of filtered polymorphic sites, suggesting that previous systemic selection had reduced genetic diversity. Despite this reduction, BactSys responded more strongly to subsequent oral selection, indicating that the slower adaptation of ContSys is unlikely to be explained by lower levels of overall genetic diversity. Instead, our results suggest that previous selection shaped the composition and frequencies of genetic variants available for subsequent adaptation, thereby influencing the evolutionary response to a new selective pressure.

### Adaptation involved multiple components of the infection outcome

Experimental evolution studies in *Drosophila* have shown that adaptation to infection can be strongly route-specific. Populations selected under one pathogen or route do not necessarily gain protection under another, and improved survival is not always accompanied by obvious costs in other traits (Mondotte & Saleh, 2018). These previous studies suggest that systemic and oral infection engage partly overlapping defense processes. Our results fit this view: bacterial clearance changed in some sex- and generation-specific cases, but it did not fully mirror the survival trajectories. Thus, adaptation to oral infection likely shares bacterial-load control mechanisms with systemic infection responses, which involves a variety of components, such as tolerance, epithelial recovery, or gut physiological resilience.

Bacterial clearance measures one component of infection outcome related to resistance, because it captures how rapidly bacterial burden declines. In females, BactSysOral flies cleared bacteria faster than ContSysOral flies early in selection, when they were still recovering from an initially lower survival level. Faster clearance may therefore have contributed to the steeper early improvement of BactSysOral females. Later, however, BactSysOral females survived better than ContSysOral females without a detectable clearance advantage. This later survival advantage is more consistent with other components of infection outcome, such as tolerance, in relation to damage control, epithelial recovery, or physiological resilience, acting in addition to bacterial-load reduction (Akyaw *et al*., 2025).

Males followed a related but distinct pattern. Early in selection, BactSysOral and ContSysOral males did not differ in bacterial clearance, even though the segmented survival model still supported a two-phase adaptive response. Later, BactSysOral males both survived better and cleared bacteria faster than ContSysOral males, suggesting that improved pathogen control became a more visible component of the male response. However, our clearance assays cannot distinguish bacterial killing from changes in gut transit or defecation, and they do not exclude tolerance-like mechanisms. This is important because recent work on oral *P. entomophila* infection shows that sex differences in survival outcomes can arise from gut physiology as much as from bacterial burden itself. Rubinic et al. (Rubinić *et al*., 2025) found that females are more vulnerable to pathogen-induced oxidative stress, defecation blockage, pathogen persistence, and gut damage, whereas males show stronger metabolic and antioxidant responses that help maintain intestinal transit and clear the pathogen. In addition, given the adaptive history of BactSysOral, our observation of faster bacterial clearance (especially in females) is consistent with reports that *P. entomophila* oral infection triggers a strong systemic response due to translocation of peptidoglycan fragments produced by the bacteria while in the gut (Vodovar *et al*., 2005; Westlake, Hanson & Lemaitre, 2024).

Overall, our results are consistent with the notion that, despite a common evolutionary history, females and males rely differently on a combination of resistance, tolerance, and gut physiological resilience, both at the physiological and adaptive levels. Different mechanisms appear to have contributed at different stages of experimental evolution, suggesting that the genetic architecture of adaptation involved multiple physiological components with distinct evolutionary dynamics.

### Oral adaptation changed systemic performance without erasing history

In our previous work with the ancestral populations, systemic infection with *P. entomophila* produced a clear increased survival under challenge (Martins et al., 2013). If oral selection had simply produced a generalized improvement in immunity, we would expect systemic performance to remain high or improve further. Instead, BactSys-derived populations retained higher systemic-infection survival than ContSys-derived populations, but oral-selected populations showed reduced systemic performance relative to their corresponding ancestral histories. Therefore, the legacy of systemic selection imposed an adaptive cost to the oral adaptation.

This pattern helps reconcile two ideas that can otherwise seem contradictory. On one hand, systemic and oral infections engage different barriers, tissues, and immune pathways, so adaptation to one route need not transfer cleanly to another (Buchon *et al*., 2009; Broderick & Lemaitre, 2012; Martins *et al*., 2013). On the other hand, route specificity does not imply that adaptation to a new infection route occurs independently of previous evolutionary history. Although the two routes involve distinct mechanisms, they also share components of the host immune response, providing a potential basis for evolutionary interactions between them. Consistent with this, the effects of previous systemic selection remained detectable under systemic challenge after oral selection and also influenced the subsequent response to oral infection. Thus, the consequences of past selection extended beyond the route in which it originally occurred, shaping both the trajectory of subsequent adaptation and the trade-offs associated with it.

However, the balance between these two components of shared and separate mechanisms of immune defense, may be particularly relevant in ecologically heterogeneous environments, where infection routes are unlikely to remain constant through time. Oral exposure is considered an important natural route of bacterial infection in *Drosophila*, whereas systemic infection may occur when pathogens gain access through breaches of the cuticle, for example following wounding or interactions with parasites (Vodovar *et al*., 2005; Keebaugh & Schlenke, 2014; Subasi *et al*., 2024). Thus, populations that have previously adapted to systemic infection may subsequently experience strong selection imposed by oral pathogens without systemic infection necessarily disappearing from their environment. In our experiment, adaptation to oral infection improved performance in the new selective context but partially compromised performance in the previously selected one, while preserving the relative systemic advantage of the BactSys history. This suggests that, when infection routes vary through time or space, the consequences of adaptation may depend not only on the current route of infection but also on the sequence of infection environments previously encountered. Such route-specific benefits and trade-offs could contribute to maintaining variation in host defence in natural populations (McKean & Nunney, 2008; Faria *et al*., 2015; Hulse & Bruns, 2024; Singh *et al*., 2025).

### No consistent fecundity trade-off accompanied improved survival

A central prediction from life-history and ecological immunology is that improved immune performance can carry reproductive costs, because resources invested in defense, repair, or stress responses may be unavailable for reproduction (Sheldon & Verhulst, 1996; Lochmiller & Deerenberg, 2000; Schmid-Hempel, 2005). In our experiment, oral infection reduced fecundity relative to sucrose controls, showing that infection itself imposed a reproductive cost. However, adaptation to oral infection did not produce a consistent difference in infection-specific fecundity between BactSysOral and ContSysOral populations, and the ancestral and relaxed-selection assays also showed no stable history-specific fecundity pattern.

Thus, improved oral-infection survival was not accompanied by a detectable fecundity consequence under our conditions. This does not rule out costs in other traits, environments, or life stages, and the reduced systemic-infection performance after oral selection shows that costs of adaptation were present in another infection context. However, the absence of a consistent fecundity response is important because it argues against a simple trade-off between survival and reproductive output under oral infection.

### Ecological implications of historical contingency in host defence

In natural populations, pathogen exposure is shaped by ecological interactions that determine not only which pathogens hosts encounter, but also how infection is established (Martins *et al*., 2013; Chambers *et al*., 2014; Eleftherianos, Tafesh-Edwards & Mohamed, 2022). Infection routes may vary across ecological contexts, potentially exposing populations to different selective pressures even when pathogen identity remains unchanged. Our findings suggest that such heterogeneity may have important evolutionary consequences. Adaptation to one infection route can alter subsequent responses to another, while improvements in one context may come at the expense of performance in another. These trade-offs raise the possibility that variation in exposure routes across space and time could contribute to maintaining diversity in host defence, consistent with broader arguments linking environmentally heterogeneous selection to the maintenance of genetic polymorphism (Laine & Tellier, 2008; Martins *et al*., 2013; Arnold *et al*., 2026).

Beyond their consequences for host adaptation, these history-dependent responses may also have implications for host–pathogen interactions at the ecological level. Our results highlight the multidimensional nature of host defence, with survival and bacterial clearance showing distinct evolutionary responses depending on sex and selection history. Such differences could potentially influence pathogen persistence within hosts and, ultimately, patterns of transmission. Indeed, variation in bacterial shedding has been identified as a relevant component of infection heterogeneity in *Drosophila* (Siva-Jothy *et al*., 2018). Whether evolutionary history influences these specific processes, and whether such effects translate into changes in infection prevalence or pathogen dynamics, remains to be established.

Combining experimental evolution with variation in infection routes, and extending measurements beyond individual survival to include pathogen shedding, transmission, and population-level fitness, could help establish when historical contingency has ecological consequences beyond the evolutionary trajectories of individual host populations. More broadly, our findings suggest that predicting host adaptation requires considering not only the pathogen and current selective environment, but also the evolutionary history of the population. An important direction for future research will therefore be to examine how the sequence, frequency, and ecological context of pathogen exposure shape evolutionary responses and their consequences for host–pathogen dynamics (Silva & King, 2025).

## AUTHOR CONTRIBUTIONS

**Priscilla A. Akyaw:** Data curation, Formal analysis, Investigation, Methodology, Project administration, Validation, Visualization, Writing – original draft, Writing – review & editing.

**Élio Sucena:** Conceptualization, Funding acquisition, Investigation, Methodology, Project administration, Resources, Supervision, Writing – original draft, Writing – review & editing.

**Elvira Lafuente:** Conceptualization, Data curation, Formal analysis, Investigation, Methodology, Project administration, Validation, Visualization, Writing – original draft, Writing – review & editing.

## DATA AVAILABILITY

The raw phenotypic data underlying the analyses are provided in the Supplementary Materials. The raw Pool-seq data will be deposited in a publicly accessible repository upon acceptance.

## ACKNOWLEDGEMENTS

We thank Diogo Roque, Sofia Nunes, Tânia F. Paulo, João Picão Osório, and Gabriele Maria Sgarlata for their help with population logistics and for insightful comments on the project along the way. We also thank Vítor G. Faria and Nelson E. Martins for their *historical role* in this project.

