## Supplementary Material for "Historical contingency is revealed by trade-offs upon sequential adaptation to different infection routes"

### Supplementary Figures

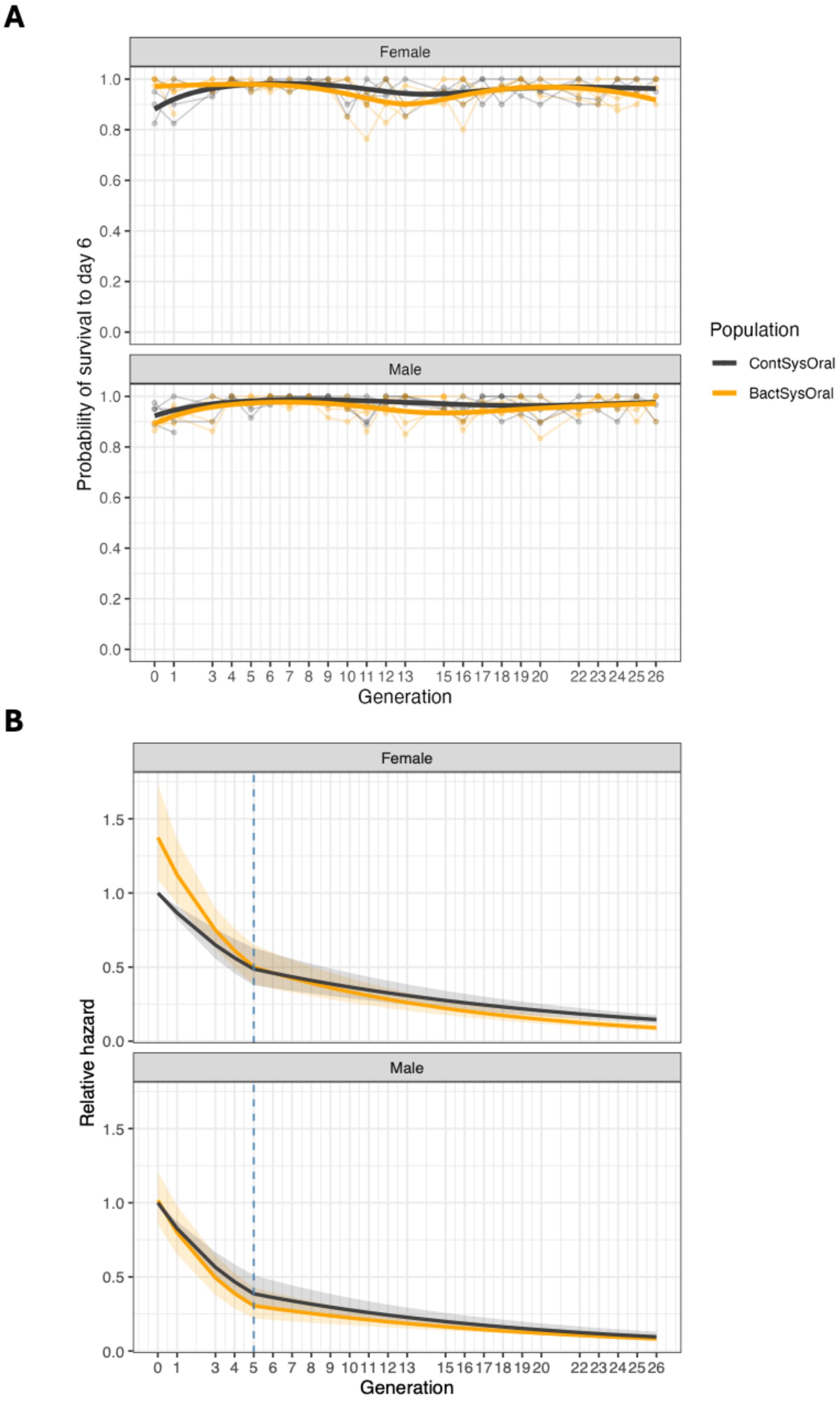

**Figure S1. Survival under sucrose control treatment across the selected trajectory.**

Probability of survival to six days after sucrose control exposure across generations. Replicate populations are shown as thin lines and model-predicted population means as thick lines. Survival under sucrose remained high throughout the experiment, indicating that the directional survival response observed under PeWT was infection-specific. Gaps at G2, G14, and G21 indicate generations in which survival was not assayed because of technical constraints.

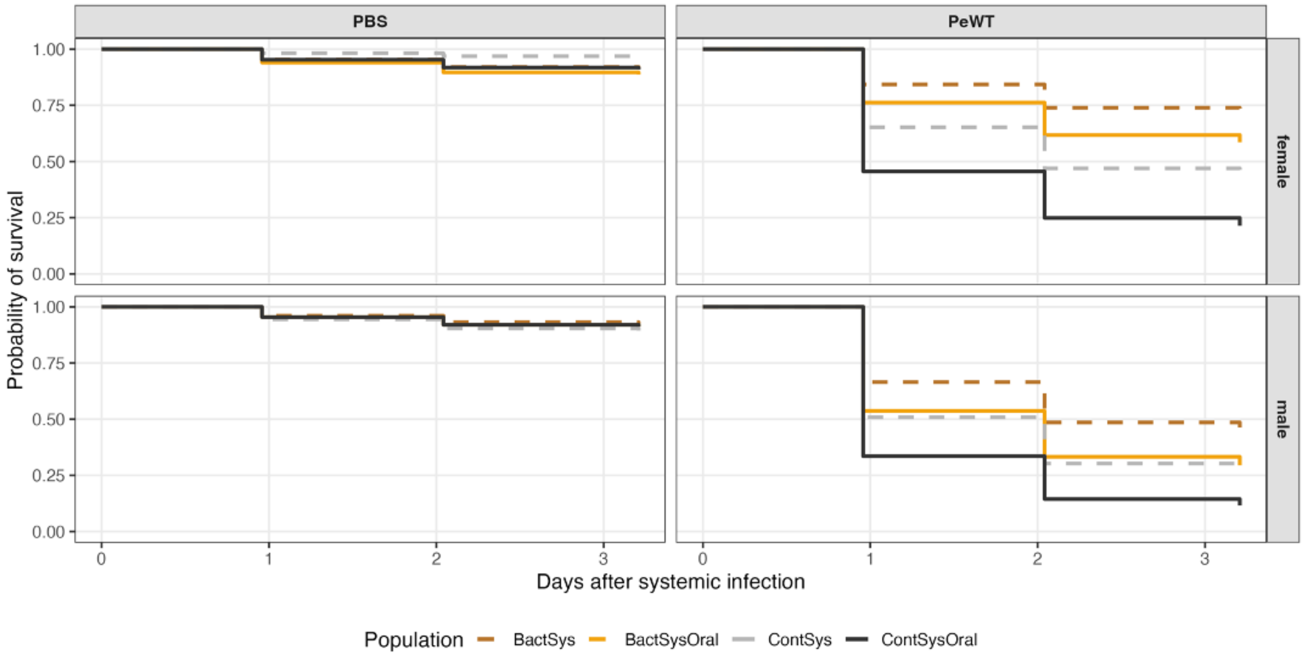


**Figure S2. Survival dynamics after systemic challenge in ancestral and after-selection populations.**

Survival curves after systemic challenge with control PBS or *Pseudomonas entomophila* (PeWT), shown separately for females and males. Dashed lines show ancestral BactSys and ContSys populations, and solid lines show after-selection BactSysOral and ContSysOral populations. BactSys-derived populations are shown in orange tones and ContSys-derived populations in grey/black tones. PBS-pricked controls showed high survival across populations, whereas PeWT challenge revealed higher systemic-infection survival in BactSys-derived histories than in ContSys-derived histories.

### Supplementary Tables

**Table S1. Generation-specific oral-survival model contrasts.**

BactSysOral versus ContSysOral hazard-ratio contrasts from Cox mixed-effects models, evaluated at each generation in which survival was assayed and reported separately by sex and treatment. Hazard ratios below 1 indicate lower mortality in BactSysOral than in ContSysOral. The table provides log hazard ratios, standard errors, test statistics, *p*-values, hazard ratios, and 95% confidence intervals.

**Table S2. Complete bacterial-clearance statistics.**

Population contrasts from analyses of bacterial clearance following oral *Pseudomonas entomophila* infection, reported by generation and sex. The table includes contrasts from the raw dilution-count model and the two-part analysis separating the probability of zero detectable colony-forming units from bacterial-load dynamics among flies with detectable bacteria. Estimates, standard errors, test statistics, *p*-values, 95% confidence intervals, and estimated bacterial half-lives are provided where applicable.

**Table S3. Genomic analyses of the G0 populations.**

Summary of Pool-seq analyses comparing BactSys and ContSys populations at the beginning of oral selection. The table contains the G0 sample map, pairwise differentiation estimates, sensitivity of Hudson-type $F_{ST}$ estimates to assumed pool sizes of 120, 125, and 130 flies, expected heterozygosity estimates for each pool, and the population-level comparison of heterozygosity. Heterozygosity was estimated across the common filtered set of polymorphic sites and should not be interpreted as absolute genome-wide nucleotide diversity.

**Table S4. Individual-level oral-survival data.**

Individual survival records for BactSysOral and ContSysOral populations across the oral-selection experiment. The dataset includes generation, population, replicate population, treatment, sex, vial identity, observation time, survival status, and event indicator. Individuals alive at the final observation were retained as right-censored observations. Survival was not assayed at G2, G14, or G21 because of technical constraints.

**Table S5. Bacterial-clearance data.**

Colony-count data obtained following oral infection with *Pseudomonas entomophila*. The dataset includes generation, population, replicate population, sex, sampling time, fly and plate identifiers, dilution, colony count, dilution-adjusted bacterial-load measures, and associated experimental information. Repeated dilution measurements from the same fly are identified by the corresponding fly-level identifier.

**Table S6. Individual-level systemic-survival data.**

Individual survival records from systemic-infection assays comparing ancestral BactSys and ContSys populations with BactSysOral and ContSysOral populations after oral selection. The dataset includes population, evolutionary stage, selection history, treatment, replicate population, sex, vial identity, observation time, survival status, and event indicator. Individuals remaining alive at the final mortality check were retained as right-censored observations.

**Table S7. Fecundity data.**

Daily fecundity and survival records from assays conducted during oral selection and from the ancestral and relaxed-selection assays. The dataset includes population, replicate population, generation or evolutionary stage, treatment, sex, vial identity, daily pupal counts, female survival information, and derived daily fecundity per surviving female. These records were used to calculate vial-level fecundity and fecundity under *Pseudomonas entomophila* infection relative to the matched sucrose control.
